# Seasonal temperature extremes drive shifts in reproductive strategy in *Cassiopea xamachana*

**DOI:** 10.64898/2026.10.09.758128

**Authors:** Mikayla M. Clark, Marie E. Strader

## Abstract

Large observable densities of jellyfish (Scyphozoa) can seemingly appear out of nowhere but can often be explained by life-history transitions triggered by temperature and host associations with microorganisms. Using a globally distributed scyphozoan, we demonstrate that seasonal temperature extremes trigger polyps to differentially allocate effort in growth and reproductive output and strategy mostly independent of their microbiomes. *Cassiopea xamachana* polyps were reared under winter (20°C), ambient (28°C), and summer (33°C) temperatures with either antibiotic-treated (ABS) or microbe-exposed (MES) communities. At 20°C, polyps grew and reproduced through budding but did not strobilate. At 28°C, polyps sustained the greatest ephyra production, whereas 33°C accelerated first ephyra release but subsequently truncated strobilation and yielded smaller ephyrae with poorer condition and survival. These patterns indicate that plasticity induced by seasonal temperature extremes may optimize differential allocation towards asexual and sexual propagation. Microbial depletion restructured polyp-associated communities, yet only minimally altered growth and reproduction. Taken together, *C. xamachana* polyps are likely to overwinter, where they maximize growth and asexual reproduction. As temperatures warm, it triggers a shift in resource allocation towards strobilation. However, the warmest summer temperatures reduced polyp and ephyra fitness, with the magnitude of effect modified by their microbiome.

## Introduction

Temperature is an important environmental driver of host physiology, and in many taxa, life-history strategies strongly covary with seasonal thermal conditions through shifts in host metabolic demand and energy allocation towards growth, development, or reproduction [1–3]. This capacity to modify life-history strategies in response to environmental conditions represents a form of phenotypic plasticity that may be particularly important in seasonally variable environments (reviewed in [4–6]). Similarly, host-associated microbial communities are sensitive to temperature, with changes in thermal conditions altering growth rates, metabolic activity, abundance, and community composition [7–9]. Marine thermal environments are rapidly shifting through both increasing average temperatures and the frequency, duration, and intensity of extreme thermal events [10–13]. Consequently, host responses to changing thermal environments likely reflect the combined effects of temperature on host physiology and the temperature-associated changes in the composition and function of their associated microbial communities.

Microorganisms can contribute to host performance through nutrient acquisition and metabolism, with holobiont energetics emerging from the metabolic integration of hosts and their microbial partners [14–16].Because metabolic processes rely on coordinated exchanges among members of the holobiont, changes in temperature can alter nutrient cycling and carbon transfer, with downstream consequences for host growth, reproduction, and survival [16–18]. Microbial contributions to host metabolism may translate directly to variation in fitness-related traits as growth, reproduction, and development are energetically demanding processes. In the moon jellyfish, *Aurelia* spp., disruption or absence of the native microbiome reduced asexual budding and impaired the asexual polyp-to-medusa transition (strobilation) [19–21]. Likewise, microbial depletion inhibited asexual budding in the common hydra, *Hydra vulgaris*, an effect that was reversed following microbiome reestablishment [22]. Microbial biofilms and bacterial metabolites also provide settlement and metamorphic cues to larvae, including cnidarians [23], annelids [24], and bivalves [25], highlighting the role of host-associated microbes in regulating developmental transitions [26]. Together with their effects on host metabolism, these microbial influences may ultimately shape large-scale population dynamics.

Upside-down jellyfish, *Cassiopea* spp., are common, globally distributed photosymbiotic scyphozoans that can form blooms, linking them as potential indicators of environmental change [27,28]. Microbial partners regulate early life-history transitions in *Cassiopea* spp. Bacterial cues induce larval settlement and metamorphosis into the benthic polyp stage [29,30], which continually reproduce asexually through budding and strobilation [31]. Strobilation, which produces free-swimming ephyrae that develop into sexually mature adults, is dependent upon the establishment of symbiosis with photosynthetic dinoflagellates of the family Symbiodiniaceae [32–34].

Establishment of this algal symbiosis also restructures the associated bacterial community, with several taxa becoming enriched during strobilation, highlighting the interconnected roles of the host, algal symbionts, and microbiome in regulating *Cassiopea* development [35]. Because the *Cassiopea* life cycle is regulated by both environmental and microbial cues, this system provides a valuable opportunity to disentangle how microbial communities influence host responses under various thermal regimes.

## Materials and Methods

### Experimental Desig

*Cassiopea xamachana* polyps were sourced from Key Largo, Florida in 2022 and maintained in laboratory culture at 28°C. Polyps (N = 900 total) were randomly assigned to 200 mL beakers (18 beakers; n = 50 polyps/beaker), and beakers were randomly assigned to one of six treatment conditions in triplicate, consisting of combinations of three temperatures and two microbial conditions (Fig. 1A). Temperature treatments included winter (20°C), ambient (28°C), and summer (33°C) conditions, spanning seasonal temperature extremes experienced in the wild (Fig. S1). Microbial conditions consisted of artificial seawater with undisturbed microbial flora (microbe-exposed; MES) or depleted microbial flora (antibiotic-treated, sterile-filtered; ABS). MES was prepared by mixing Instant Ocean Sea Salt with RO water to 28 parts per thousand (ppt) and circulating it through the existing laboratory reservoir system, which had initially been established using Dr. Tim’s Aquatics Nitrifying Bacteria and Seachem Stability ® Water Conditioner. ABS was generated by sterile-filtering MES through a 0.22 µm filter and adding an antibiotic cocktail of 100 mg/L neomycin, 130 mg/L streptomycin, and 100 mg/L penicillin.

**Figure 1.**
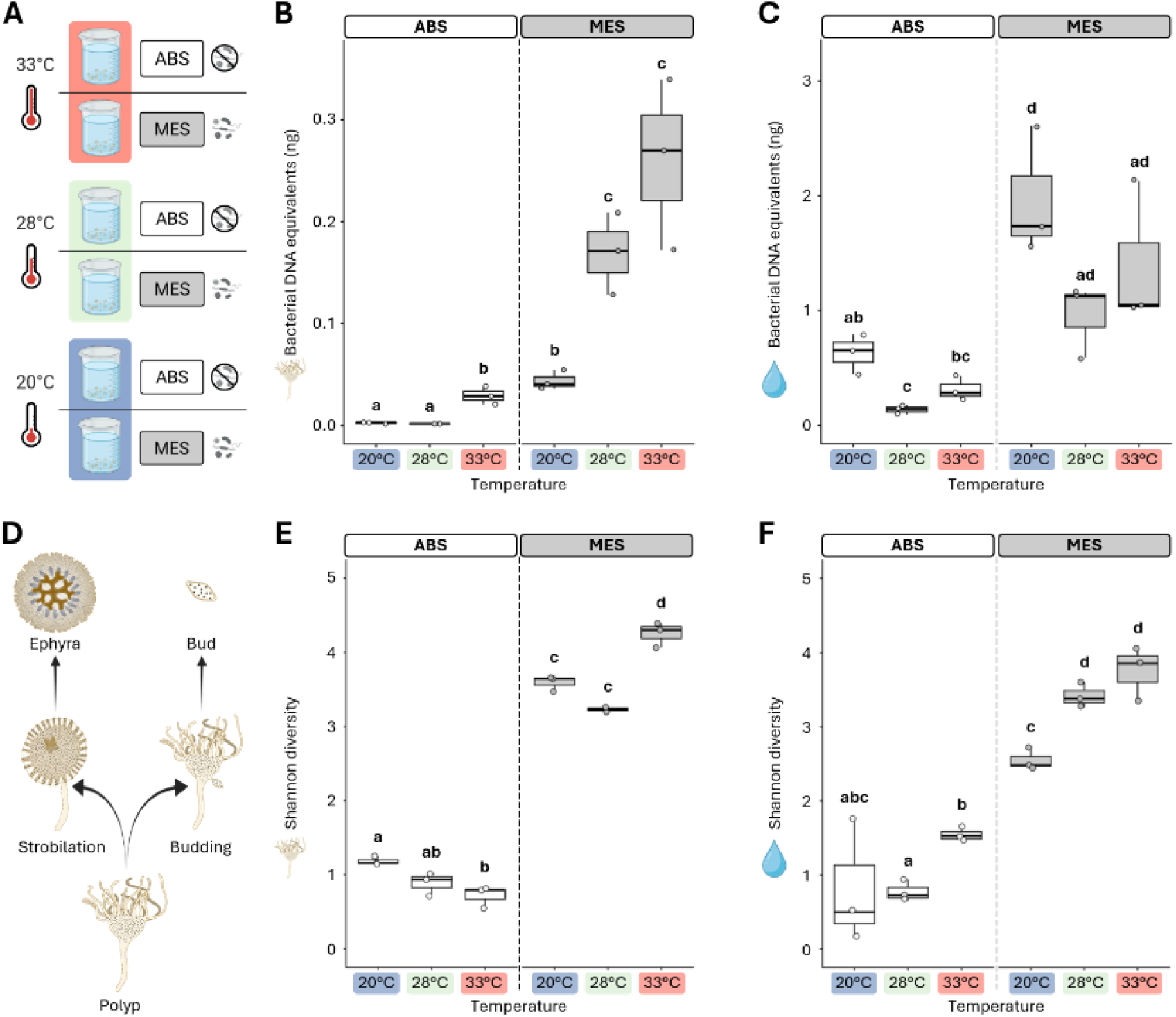
Antibiotic treatment reduces microbial presence and diversity in both polyp microbiomes and surrounding water. (A) Experimental design consisting of three temperatures (20°C, 28°C, 33°C) and two microbial treatments: antibiotic-treated sterile-filtered artificial seawater (ABS) and microbe-exposed artificial seawater (MES). (B-C) 16S bacterial DNA equivalents (ng), a proxy for bacterial load, in polyp (B) and water (C) samples at T_35_. (D) Alternative modes of asexual reproduction in *Cassiopea* polyps. Budding produces buds (n ≥ 1 offspring per event), whereas strobilation produces ephyrae (n = 1 offspring per event). (E-F) Shannon alpha diversity of polyp-associated (E) and water-associated (F) bacterial communities at T_35_. Boxes represent the median and interquartile range (IQR), with whiskers extending to extremes within 1.5× the IQR. Points represent biological replicates. Letters denote significant differences among estimated marginal means (all *p*_*adj*_ ≤ 0.02). See Table S1 for all model statistics and pairwise comparisons.

For the 20°C and 33°C treatments, polyps were acclimated from 28°C to their assigned treatment temperatures over one week in water baths under a 12:12 hour light:dark photoperiod at approximately 85-100 µmol photons/m^2^, with temperatures adjusted 1-1.5°C per day. Once the treatment temperatures were reached, polyps received a complete water change to initiate the microbial treatment (T_0_). Polyps assigned to the microbially-depleted condition were rinsed and transferred into ABS water, whereas polyps assigned to the microbe-exposed condition received a complete water change with MES water. Polyps were then reared under their assigned conditions for five weeks (35 days; T_35_) in temperature-controlled incubators under the same 12:12 hour light:dark cycle. Polyps were fed *Artemia* nauplii *ad libitum* twice weekly, followed by 100% water changes at the assigned temperature and microbial treatment.

### Morphological and Reproductive Measurements

To quantify changes in polyp abundance and asexual reproduction, polyps, buds, and ephyrae were enumerated at each feeding on experimental days 0, 3, 7, 10, 14, 17, 21, 24, 28, 31, and 35. Because polyp mortality could not be reliably identified prior to complete tissue degradation, longitudinal polyp counts were used to assess net changes in abundance, reflecting both polyp loss and the development of new polyps from rapidly settled buds, rather than direct survival. During water changes, polyps were counted and returned to their respective beakers, asexually produced buds were counted and discarded, and ephyrae were counted and pooled by treatment. Ephyrae were maintained and fed twice weekly alongside polyps for the remainder of the experiment. The timing of peak bud production was defined as the census timepoint at which the maximum number of buds was recorded for each replicate, and the timing of first ephyra release was recorded as the first census timepoint at which a free-swimming ephyra was observed. To assess changes in budding associated with strobilation, mean bud production was calculated separately for each replicate across census timepoints occurring before and after the first ephyra was released. The decline in budding following the first ephyra release was calculated as the difference between mean bud production before and after that census timepoint.

Polyps were imaged to quantify calyx width at T_0_ (n=5 per replicate) and in all surviving polyps at T_35_ on a Zeiss Primostar 3 compact microscope (4X magnification) using the Zeiss ZEN blue software (version 3.7.4). Measurements were performed in ImageJ/FIJI (version 1.54p; [36]) and calibrated using a 1 mm micrometer (Edmund Optics). Ephyrae from biological replicates were pooled by treatment and imaged at collection using a TG-7 digital camera (OM System) with a standardized in-frame ruler for scale calibration. Ephyra size was quantified in ImageJ/FIJI (version 1.54p) using three diameter measurements per individual, which were averaged to obtain a single mean diameter for each ephyra. At the final census, the number of ephyrae remaining in each treatment pool was recorded, and mortality was calculated as the cumulative number produced minus the number remaining. Ephyrae present at the final census were classified as survivors and further categorized as motile if exhibiting normal pulsing behavior or non-motile if present but not pulsing.

### DNA Extractions

To quantify microbial load and characterize microbiomes, polyps were collected for DNA extraction at T_0_ and T_35_, with pooling strategies differing between sampling timepoints. At T_0_, 5 polyps from each of the three biological replicate beakers within a treatment were pooled, generating one composite sample containing 15 polyps per treatment (N = 6 samples total). At T_35_, approximately 25 polyps were collected from each biological replicate beaker and were pooled within, but not across, beakers, generating three independent samples per treatment (N = 18 samples in total). Prior to extraction, samples were centrifuged at 10,000 rpm for 3 minutes to remove residual water and immediately processed using a standard cetyltrimethylammonium bromide (CTAB) protocol. To characterize microbial communities in the surrounding water, approximately 200 mL of water was collected from each beaker at T_35_ and filtered through 0.45 µm membrane filters. DNA was extracted from the filters using a modified ZymoBIOMICS DNA Miniprep Kit (Zymo Research; D4300) protocol [37].

### Microbial Quantification

Approximate bacterial load was quantified by qPCR using a Bio-Rad CFX Opus 96 system. The TaqMan Microbe Detection Assay 16S Pan-Bacterial Control (Ba04930791_s1) and TaqPath BactoPure Microbial Detection Master Mix with ROX (ThermoFisher; A52699) were used to amplify and detect bacterial 16S rRNA gene sequences. A six-point serial dilution (0.0001 - 10 ng) of the ZymoBIOMICS Microbial Community DNA Standard (Zymo Research; D6305) was used to generate a standard curve, and molecular grade nuclease-free water served as a negative control. Sample DNA extracts were normalized to 5 ng/µL, and 2 µL of template was used per reaction. All reactions were performed in technical triplicate. C_q_ values were converted to bacterial DNA equivalents using the six-point standard curve, and relative bacterial load was expressed as bacterial DNA equivalents per 10 ng total DNA input.

### 16S Sequencing

DNA from polyp and water samples was sent to Novogene for 16S rRNA gene amplicon sequencing targeting the V3-V4 region using primer sequences CCTAYGGGRBGCASCAG and GGACTACNNGGGTATCTAAT. Libraries were sequenced on an Illumina NovaSeq 6000 platform. Raw sequence data were processed using QIIME2 (version 2025.7-Amplicon; [38]). Demultiplexed reads were quality filtered, denoised, and merged using the DADA2 plugin (q2-dada2 version 2025.7.0; [39]) to generate amplicon sequence variants (ASVs). Taxonomic assignments were performed using a naive Bayes classifier trained using the SILVA 138.2 database trimmed to the V3–V4 region [40]. Because *C. xamachana* is photosymbiotic and algal-derived sequences were expected to contribute to the dataset, we quantified the proportion of non-target ASVs relative to bacterial ASVs across all samples (Fig. S5). For all downstream bacterial community analyses, non-target sequences, including chloroplast, mitochondrial, eukaryotic, archaeal, and undetermined reads, were removed prior to analysis.

Bacterial community analyses were conducted in RStudio (version 2026.04.0+526) using the packages qiime2R (v0.99.6; [41]), phyloseq (v1.54.2; [42]), vegan (v2.7.5; [43]), and tidyverse (v2.0.0; [44]). For diversity analyses, samples were rarefied to the minimum sequencing depth of 15,530 reads per sample. For taxonomic composition and relative abundance, analyses were restricted to T_35_ polyp samples and rarefied to the minimum sequencing depth of the subset (16,997 reads per sample). Relative abundances are presented from phylum to genus (Tables S5-S9). Shannon alpha diversity [45] was calculated to assess within-sample diversity. Additional alpha diversity metrics included observed ASVs and Faith’s phylogenetic diversity (Fig. S2A-D). Beta diversity was evaluated using Bray-Curtis dissimilarity [46] and weighted UniFrac distance [47], incorporating taxon abundance and phylogenetic relatedness, respectively, and visualized using Principal Coordinates Analysis (PCoA). Additional beta diversity metrics included Jaccard dissimilarity and unweighted UniFrac distance (Figure S2E-F).

Differential abundance of bacterial taxa among the six treatment groups was assessed using Analysis of Compositions of Microbiomes with Bias Correction 2 (ANCOM-BC2 v2.12.1; [48]) on non-rarefied count data. Analyses were restricted to T_35_ polyp samples and were conducted separately at class, order, and genus levels after removing taxa present in fewer than 10% of samples. Global tests were used to identify taxa differing among treatment combinations, with pairwise contrasts used to localize differences where supported. *P*-values were adjusted for multiple comparisons using the Benjamini-Hochberg correction [49], and significant taxa were required to pass pseudo-count sensitivity analysis. Robust global results are reported in Table S4.

### Statistical Analyses

All statistical analyses were conducted in RStudio (version 2026.04.0+526). Unless otherwise specified, linear models were fit with temperature, microbial treatment, and their interaction as fixed effects (Temperature * Microbial_Treatment). Linear model assumptions were evaluated using Shapiro–Wilk tests of residual normality [50], Levene’s tests for homogeneity of variance [51], and global validation of linear model assumptions using the package gvlma (v1.0.0.3; [52]). Model diagnostic plots were also visually inspected. The significance of linear model terms was assessed using Type III Analysis of Variance (ANOVA; package car; [53]). Post-hoc pairwise comparisons were conducted using estimated marginal means (package emmeans; [54]) with Tukey-adjusted *p*-values. Compact letter displays used to visualize significant differences among treatment groups were generated from Sidak-adjusted estimated marginal means using the package multcomp [55]. For models exhibiting heteroscedasticity (Shannon diversity water samples), Type III ANOVAs were conducted using HC3 heteroscedasticity-consistent covariance estimates [56]. The same HC3 correction was applied to subsequent estimated marginal means comparisons. Polyp and water samples were analyzed separately for bacterial load and alpha diversity as they represent distinct host-associated and environmental microbial compartments.

When parametric model assumptions were not met, non-parametric tests were used. The timing of peak bud production was compared among treatment groups using a Kruskal-Wallis test, with post-hoc comparisons among temperatures conducted using Dunn’s tests with Benjamini-Hochberg-adjusted *p*-values [49,57]. The timing of peak bud production was also compared between microbial treatments using a Wilcoxon rank-sum test. The timing of first ephyra release was compared among the four strobilating treatments using a Kruskal-Wallis test, with temperature and microbial treatment effects subsequently assessed using Wilcoxon rank-sum tests.

Changes in mean bud production before and after first ephyra release were assessed using paired Wilcoxon signed-rank tests. The relationship between cumulative bud and ephyra production was assessed using Spearman’s rank correlation coefficient (ρ) to evaluate evidence of a tradeoff between the two modes of asexual reproduction.

Differences in bacterial community composition among treatment groups were assessed for T_35_ polyp samples using Permutational Multivariate Analysis of Variance (PERMANOVA) with temperature, microbial treatment, and their interaction (Temperature * Microbial_Treatment) using the package vegan [43].

PERMANOVAs were conducted using 999 permutations, with model terms evaluated sequentially. Homogeneity of multivariate dispersion was assessed among the six treatment groups using Permutational Analysis of Multivariate Dispersions (PERMDISP), also with 999 permutations (package vegan). For weighted UniFrac, a Lingoes correction was applied during PERMDISP to account for the non-Euclidean distance matrix [58].

## Results

### Antibiotic treatment depletes bacterial DNA abundance and diversity in both host and surrounding water

To quantify microbiome depletion, we used qPCR to estimate bacterial DNA abundance and 16S sequencing to characterize microbial diversity. Because 16S gene copy number varies among bacterial taxa [59], qPCR results are reported as bacterial DNA equivalents detected from a known amount of template DNA, which serves as a proxy for bacterial load. In polyp samples, bacterial DNA equivalents differed significantly by temperature, microbial treatment, and their interaction (all *p*_*ANOVA*_ < 0.001; Fig. 1B; Table S1). Within ABS polyps, bacterial DNA equivalents were significantly greater at 33°C than at both 20°C and 28°C (both *p*_*adj*_ < 0.001), which did not differ significantly (*p*_*adj*_ = 0.724). Within MES polyps, bacterial DNA equivalents were significantly lower at 20°C than at both 28°C and 33°C (both *p*_*adj*_ ≤ 0.001), which did not differ significantly (*p*_*adj*_ = 0.507). In water samples, bacterial DNA equivalents varied significantly with temperature and microbial treatment (both *p*_*ANOVA*_ < 0.001; Fig. 1C; Table S1), but not their interaction (*p*_*ANOVA*_ = 0.187).

Antibiotic treatment reduced bacterial alpha diversity in both polyps and the surrounding water (Fig. 1E-F). In polyps, Shannon diversity differed significantly by temperature, microbial treatment, and their interaction (all *p*_*ANOVA*_ < 0.001; Fig. 1E; Table S1) and was approximately fourfold greater in MES than ABS polyps. Within MES polyps, diversity was significantly greater at 33°C than at 20°C and 28°C (both *p*_*adj*_ ≤ 0.001), whereas within ABS polyps, only 20°C and 33°C differed significantly (*p*_*adj*_ = 0.009). In water samples, Shannon diversity varied significantly with temperature (*p*_*HC3*_ = 0.006) and microbial treatment (*p*_*HC3*_ < 0.001; Fig. 1F; Table S1), but not their interaction (*p*_*HC3*_ = 0.210). Diversity was greatest in MES water at 33°C and lowest in ABS water at 20°C and 28°C.

### Temperature and bacterial load influence fitness-related traits

To assess effects of temperature and microbial treatment on host growth, we quantified changes in polyp calyx width from T_0_ to T_35_. Growth differed significantly among temperatures (*p*_*ANOVA*_ < 0.001; Fig. 2A; Table S2). Polyps at 20°C grew under both microbial treatments, reaching approximately 1.5× their initial size, whereas polyps at 28°C and 33°C declined to approximately 0.85× and 0.68× their initial size, respectively. Microbial treatment also had a significant effect on growth (*p*_*ANOVA*_ = 0.013), with ABS polyps at 28°C and 33°C approximately 26% larger at T_35_ than their MES counterparts (Fig. 2A; Table S2).

**Figure 2.**
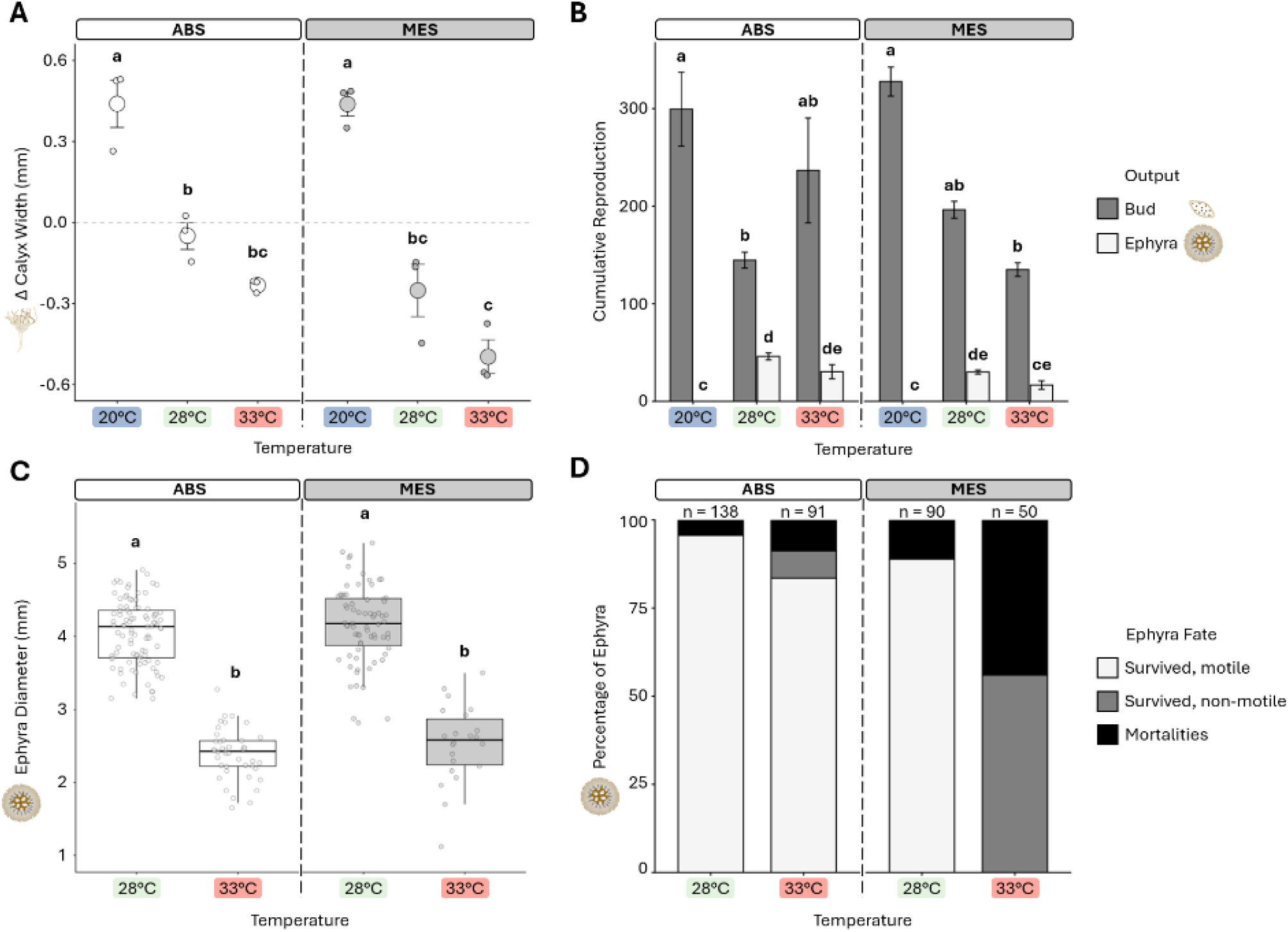
Temperature elicited stronger effects on asexual reproduction and growth than microbial depletion. (A) Change in polyp calyx width from T_0_ to T_35_. Large circles represent treatment means ± standard error, and small circles represent biological replicates. (B) Mean cumulative production of buds and ephyrae ± standard error, with each reproductive mode analyzed separately. (C) Ephyra diameter measured at collection for all ephyrae produced within each treatment. The 20°C treatments did not produce ephyrae. Boxes represent the median and interquartile range (IQR), with whiskers extending to extremes within 1.5× the IQR. Points represent individual ephyrae. (D) Ephyra fate at T_35_. Letters denote significant differences among estimated marginal means (all *p*_*adj*_ ≤ 0.02). See Table S2 for all model statistics and pairwise comparisons.

Both modes of asexual reproduction, budding and strobilation, were quantified as metrics of organismal fitness. Cumulative bud production was significantly influenced by temperature (*p*_*ANOVA*_ < 0.001; Fig. 2B; Table S2), with a significant temperature × microbial treatment interaction (*p* = 0.038), while the main effect of microbial treatment was not significant (*p* = 0.758). Bud production was highest at 20°C under both microbial treatments and lowest in 28°C ABS and 33°C MES. Cumulative ephyra production was also significantly influenced by temperature (*p*_*ANOVA*_ < 0.001; Fig. 2B; Table S2), with no ephyrae produced at 20°C. In contrast to budding, microbial treatment significantly influenced ephyra production (*p*_*ANOVA*_ = 0.008), with ABS polyps producing approximately 64% more ephyrae than MES polyps. However, neither pairwise comparisons between microbial treatments at each temperature (Table S2) nor the temperature × microbial treatment interaction (*p*_*ANOVA*_ = 0.120) were significant.

In addition to producing the greatest number of ephyrae, polyps at 28ºC also produced larger ephyrae than those at 33ºC (*p*_*ANOVA*_ < 0.001; Fig. 2C; Table S2). Ephyrae averaged 4.1 mm in diameter when collected at 28°C compared with 2.5 mm at 33°C, an approximately 40% reduction. Neither microbial treatment (*p*_*ANOVA*_ = 0.079) nor the temperature × microbial treatment interaction (*p*_*ANOVA*_ = 0.825) significantly affected ephyra diameter (Table S2). Ephyra survival and condition at T_35_ also varied among treatments (Fig. 2D; representative images in Fig. S8B). At 28°C, ≥ 88% survived in both microbial treatments, with 4.3% mortality in ABS and 11.1% in MES. In contrast, ephyra fate diverged strongly between microbial treatments at 33°C. MES had 44% mortality, with all surviving ephyrae (56%) non-motile, whereas ABS had 8.8% mortality and 7.7% non-motile survivors, with the remaining 83.5% surviving and motile. The non-motile phenotype only occurred at 33°C. Because ephyrae were pooled by treatment, survival and condition were assessed descriptively. Overall, 28°C yielded more, larger ephyrae with high survival, whereas 33°C yielded fewer, smaller ephyrae with poorer outcomes, particularly under MES conditions.

### Temperature and microbial treatment alter the timing and allocation of asexual reproduction

Because cumulative reproductive output provides only an endpoint measure of reproductive investment, temporal patterns of budding and strobilation were also examined (Fig. 3). Neither microbial treatment at 20°C produced ephyrae, and bud production increased steadily throughout the experiment (Fig. 3A), ultimately resulting in the greatest cumulative bud production (Fig. 2B). In contrast, strobilation occurred at both 28°C and 33°C, regardless of microbial treatment, and was accompanied by a decline in bud production. The timing of first ephyra release differed significantly among the four strobilating treatments (Kruskal-Wallis, *p* = 0.036; Fig. 3B; Table S3). Ephyrae were released significantly earlier at 33°C than at 28°C (Wilcoxon rank-sum test, *p* = 0.008), whereas timing did not significantly differ between microbial treatments (*p* = 0.523).

**Figure 3.**
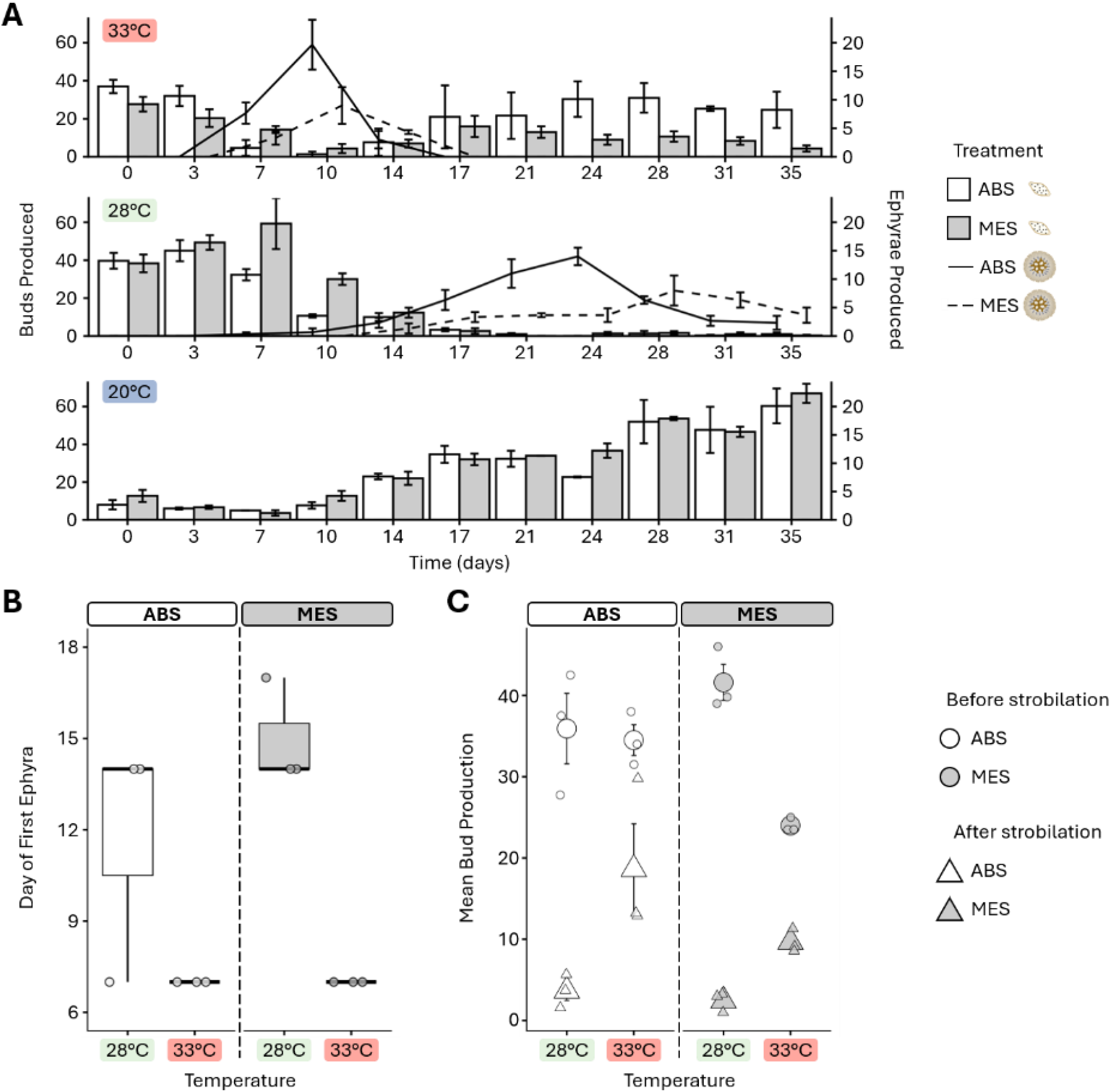
Temperature influences temporal shifts in reproductive allocation between budding and strobilation. (A) Mean bud and ephyra production per treatment across the experiment ± standard error. Bars correspond to the lefthand y-axis and represent mean buds produced per timepoint, while lines correspond to the righthand y-axis and represent mean ephyrae produced per timepoint. (B) Day of the first ephyra release. The 20°C treatments did not produce ephyrae. Boxes represent the median and interquartile range (IQR), with whiskers extending to extremes within 1.5× the IQR. Points represent biological replicates. (C) Mean bud production before (circles) and after (triangles) first ephyra release. Large points represent treatment means ± standard error, and small points represent biological replicates. See Table S3 for all model statistics and pairwise comparisons.

Budding dynamics showed a similar temperature-dependent pattern, with the timing of peak bud production differing significantly among the six treatment groups (Kruskal-Wallis, *p* = 0.019; Fig. S4A; Table S3). When grouped by temperature, peak budding occurred significantly earlier at 28°C and 33°C than at 20°C (*p*_*adj*_ = 0.007 and 0.001, respectively; Table S3) but did not differ significantly between 28°C and 33°C (*p* = 0.202). Microbial treatment did not significantly affect the timing of peak bud production (Wilcoxon rank-sum test, *p* = 0.717). The magnitude of peak bud production also differed significantly with temperature (*p*_*ANOVA*_ = 0.006; Fig. S4B; Table S3), while neither microbial treatment nor the temperature × microbial treatment interaction had a significant effect (*p*_*ANOVA*_ = 0.593 and 0.108, respectively). Polyp abundance changed over time, particularly at 33°C (Fig. S3A), but normalizing reproduction by polyp abundance did not qualitatively alter the temporal patterns (Fig. S3A-B).

Given the opposing patterns of budding and strobilation, we examined whether bud production changed after first ephyra release. Mean bud production was significantly lower after first ephyra release across all strobilating treatments (paired Wilcoxon signed-rank test, *p* < 0.001; Fig. 3C; Table S3), and remained significantly lower when 28°C and 33°C were analyzed separately (both *p* = 0.031). The difference in mean bud production before and after first ephyra release was significantly greater at 28°C than at 33°C (*p*_*ANOVA*_ < 0.001), declining by 35.7 buds at 28°C compared with 15.1 buds at 33°C. Neither microbial treatment nor the temperature × microbial treatment interaction significantly affected this decline (*p*_*ANOVA*_ = 0.409 and 0.208, respectively). Cumulative bud and ephyra production were also negatively correlated across the four strobilating treatments (Spearman’s ρ = -0.664, *p* = 0.003). Collectively, these results demonstrate a temperature-dependent shift in asexual reproductive allocation, with budding declining after first ephyra release and the magnitude of this decline differing between temperatures but not microbial treatments.

### Temperature modulates antibiotic-induced restructuring of the polyp-associated bacterial community

To determine how temperature and microbial treatment influenced polyp-associated bacterial community composition, beta diversity was assessed using abundance- and phylogeny-based dissimilarity metrics. Principal Coordinates Analysis (PCoA) of Bray-Curtis dissimilarities revealed strong separation between ABS and MES communities along axis 1, with additional clustering by temperature along axis 2 (Fig. 4A). PERMANOVA confirmed significant effects of microbial treatment (R^2^ = 0.698, *p* = 0.001), temperature (R^2^ = 0.128, *p* = 0.001), and their interaction (R^2^ = 0.128, *p* = 0.001; Table S4). Weighted UniFrac produced similar patterns (Fig. 4B), with PERMANOVA again detecting significant effects of microbial treatment (R^2^ = 0.851, *p* = 0.001), temperature (R^2^ = 0.071, *p* = 0.001), and their interaction (R^2^ = 0.054, *p* = 0.002; Table S4). Microbial treatment explained the greatest proportion of variation in abundance-weighted phylogenetic composition. PERMDISP detected no significant differences in within-group dispersion for either Bray-Curtis (*p* = 0.260) or weighted UniFrac (*p* = 0.434).

**Figure 4.**
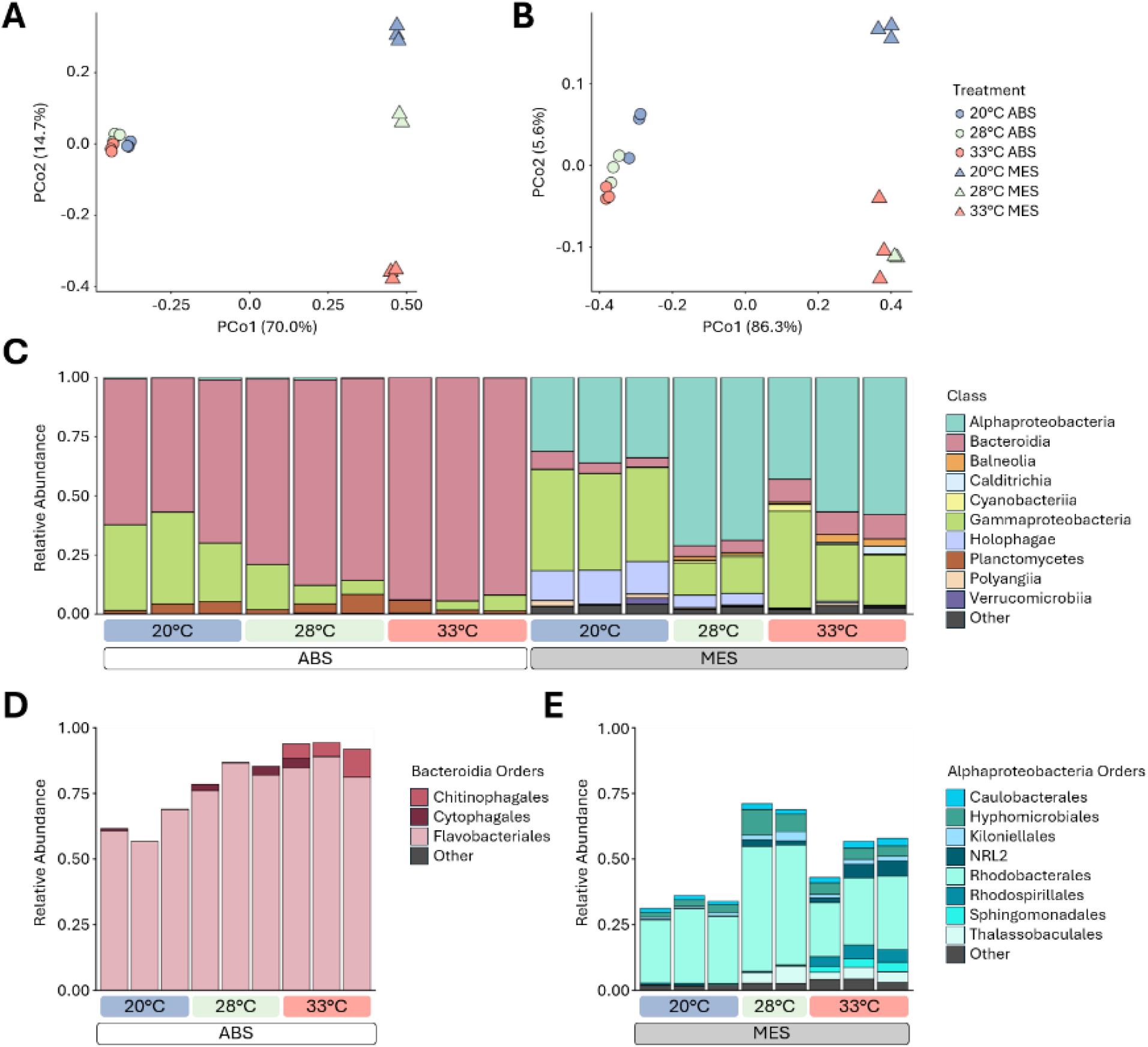
Temperature and microbial treatment altered polyp-associated bacterial community composition. (A-B) Principal Coordinates Analysis (PCoA) of beta diversity using Bray-Curtis dissimilarity (A) and weighted UniFrac distance (B). (C) Relative abundance of bacterial classes across treatments. (D-E) Relative abundance of orders within Class Bacteroidia (D), the most abundant class in ABS polyps, and Class Alphaproteobacteria (E), the most abundant class in MES polyps. The “Other” category represents unclassified bacterial taxa and taxa below the 2% total abundance threshold. See Table S4 for all model statistics and pairwise comparisons.

ABS communities were dominated by Bacteroidia, whereas MES communities contained greater proportions of Alphaproteobacteria (Fig. 4C). Alphaproteobacteria never exceeded 1% relative abundance in any ABS samples, while Bacteroidia remained below 10% across MES samples (Fig. S7B-C). Within ABS communities, Bacteroidia was predominantly order Flavobacteriales (Fig. 4D), with the genus *Muricauda* comprising approximately 62%, 81%, and 85% of the total bacterial community at 20°C, 28°C, and 33°C, respectively (Fig. S7; Table S4). In contrast, Alphaproteobacteria in MES communities were distributed across eight orders, with Rhodobacterales being the most abundant (Fig. 4E). Among reads classified to genus level, *Ruegeria*, a member of Rhodobacterales, was the most abundant in MES communities, comprising approximately 20%, 41%, and 11% of the total bacterial community at 20°C, 28°C, and 33°C, respectively (Fig. S7; Table S4). Thus, microbial treatments were characterized by distinct dominant bacterial lineages, with additional temperature-associated variation in relative abundance.

ANCOM-BC2 global tests identified 8 classes, 16 orders, and 18 genera that differed significantly among the six treatment combinations (Table S4). These included *Muricauda* (*q* = 0.001) and *Ruegeria* (*q* = 0.006), consistent with the compositional shifts described above. Pairwise differences were limited after multiple-testing correction. Alphaproteobacteria was significantly more abundant in MES than ABS at 20°C, 28°C, and 33°C (all *q* ≤ 0.009), while no order- or genus-level pairwise contrasts were significant. Together, these results demonstrate that microbial treatment restructured the polyp-associated bacterial community, while temperature further altered community composition and the relative abundance of dominant bacterial taxa.

## Discussion

### Temperature-dependent reproductive allocation may shape seasonal population dynamics

Temperature is a major environmental cue regulating asexual reproduction in scyphozoan polyps, influencing the timing and magnitude of budding and strobilation [60,61]. Here, temperature reorganized developmental and reproductive trajectories in *C. xamachana* polyps rather than uniformly suppressing or enhancing asexual reproduction. At 20°C, polyps reproduced exclusively through budding, whereas at 28°C and 33°C, they transitioned from budding to strobilation at different rates, accompanied by a pronounced decline in budding (Fig. 3). Following strobilation, reproductive trajectories diverged by temperature, with budding declining to near-zero levels alongside sustained ephyra production at 28°C, whereas at 33°C, strobilation occurred earlier, was confined to a 10-day window, and then ceased as budding resumed. These reciprocal patterns suggest that initiation and persistence of budding and strobilation represent alternative reproductive states that are strongly temperature dependent. Similar cycling has been documented in *C. andromeda*, where polyps either returned to budding or entered another strobilation cycle following ephyra release [62,63]. Strobilation involves extensive morphological reorganization and subsequent regeneration of the feeding structures [34].

Studies in other scyphozoans indicate that food availability can influence strobilation [64] and that temperature can shift allocation between somatic growth and asexual reproduction [65]. Consistent with this tradeoff, mean calyx width increased at 20°C, where polyps were not strobilating, and decreased at 28°C and 33°C, where the calyx contributed to ephyra formation and required subsequent regeneration (Fig. 2A). However, this pattern could also reflect differences in population size structure. Although buds were removed at each census, temperature-induced rapid settlement between censuses (< 3 days), which was most prominent at 33°C, could have increased the representation of smaller, newly settled polyps and thereby contributed to the reduction in mean calyx width (Fig. S3A). Thus, differences in mean calyx width may reflect the consequences of strobilation, the presence of newly settled polyps, or a combination of both processes.

The temperature-dependent reproductive trajectories observed here provide a potential mechanism linking seasonal conditions experienced by polyps to changes in *Cassiopea* medusa populations. In the Florida Keys, Fitt & Costley [66] observed strong seasonality in *C. xamachana*, with medusae and newly strobilated ephyrae present during warmer months but absent between late November and early June. This pattern aligns with our observation that strobilation occurred at 28°C and 33°C but not at 20°C. Importantly, however, polyps at 20°C remained active, growing and reproducing through budding. Because polyp abundance reflects sexual recruitment, asexual propagation, and mortality [67], continued budding under cooler conditions could increase the number of polyps available to strobilate as temperatures rise. Field observations in *Aurelia labiata* similarly demonstrate substantial interannual variation in the timing and magnitude of strobilation despite relatively stable polyp densities [68].

Seasonal surveys of *Cassiopea* in a Caribbean mangrove also documented substantial variation in medusa density and size structure, indicating that population responses vary with environmental conditions rather than following a fixed seasonal pattern [69]. Although the seasonal abundance and fate of wild *Cassiopea* polyps remain poorly resolved, our results identify temperature-dependent reproductive transitions in polyps as one mechanism that could contribute to changes in medusa demography.

Warmer temperatures can accelerate developmental processes in scyphozoans, including the transition from polyp to ephyra [70–72]. This is particularly relevant for *C. xamachana*, which experiences temperatures near and above 33°C during summer in South Florida (Fig. S1) [66]. In our experiment, initial ephyra release occurred approximately one week earlier at 33°C than at 28°C, demonstrating accelerated development under elevated temperature (Fig. 3B). However, earlier release did not translate into sustained production or improved performance, as strobilation at 33°C subsequently ceased and ephyrae were smaller with poorer survival than at 28°C (Fig. 3A; Fig. 2C-D). A similar distinction between accelerated development and survival has been observed in *Chrysaora pacifica*, where higher temperatures produced the greatest ephyra growth but lowest survival [73]. The cessation of strobilation at 33°C may therefore reflect declining polyp condition during prolonged exposure rather than an inability to initiate reproductive development at this temperature. Although algal symbiont abundance was not quantified, polyps at 33°C exhibited conspicuous loss of pigmentation (Fig. S8), consistent with disruption of the photosymbiotic state. Because Symbiodiniaceae symbiosis is required for strobilation in *Cassiopea* [32–34], impairment of this association could have limited subsequent strobilation cycles even as budding resumed. If strobilation occurs within a bounded thermal window, changes in the timing and duration of seasonal temperatures could alter both the phenology and magnitude of ephyra production. Rather than uniformly increasing reproductive output, sustained exposure to upper seasonal temperatures may truncate this favorable window, reducing the number and quality of ephyrae entering the water column. This may become increasingly important as South Florida waters continue to warm and experience more frequent or prolonged heat anomalies [74,75].

### Microbial communities modify the consequences of temperature

Host-associated microbial communities are responsive to temperature and can, in turn, influence host physiology and thermotolerance, thereby shaping holobiont performance [76,77]. In our study, substantial reductions in bacterial abundance and diversity did not disrupt the major temperature-dependent developmental and reproductive responses of *C. xamachana* polyps but instead modified their magnitude and outcome. Across temperatures, microbial treatment significantly affected polyp calyx growth, with greater mean growth under ABS conditions. ABS polyps also produced more ephyrae than MES polyps across temperatures where strobilation occurred. Because antibiotic treatment simultaneously reduced bacterial abundance and diversity and restructured community composition, these differences cannot be attributed to bacterial depletion alone or the presence or absence of particular taxa. Instead, they reflect reproduction and growth under distinct microbial community states. The divergence between microbial states was most evident at 33°C, where ephyra condition and survival were markedly poorer under MES than ABS conditions, suggesting that microbial context modified responses to the upper seasonal temperature. Nevertheless, substantial depletion and restructuring of the polyp microbiome did not prevent budding, bud settlement, or strobilation. This persistence is notable because bacterial cues directly induce an earlier life-history transition in *Cassiopea*, namely planula settlement and metamorphosis into polyps [29,30], suggesting that dependence on bacterial cues may differ among life-history transitions.

Microbial communities can modify host responses to elevated temperature, although the direction of these effects varies among host systems. In *A. aurita* polyps, acute increases in temperature or salinity reduced survival, growth, feeding, and asexual reproduction, with stronger effects observed in bacteria-depleted polyps [78].

Similarly, antibiotic disruption of the bacterial community in *Pocillopora* corals amplified host and algal symbiont transcriptional responses associated with stress during acute heat exposure [79]. Conversely, inoculation with a selected bacterial consortium improved survival and recovery following thermal exposure in *Mussismilia hispida* corals [80]. The better condition and survival of ABS ephyrae at 33°C should, therefore, not be interpreted as evidence that bacterial depletion was inherently beneficial. Together with contrasting outcomes reported in other cnidarians, our results highlight the importance of microbial context in shaping host responses across temperatures.

### Changes in bacterial communities may influence holobiont interactions

*Cassiopea xamachana* is increasingly recognized as a model for tripartite interactions among the animal host, its Symbiodiniaceae photosymbionts, and associated bacterial communities (reviewed by [81]) [82]. Although bacterial communities of wild larvae and medusae have been characterized [82,83], the polyp-associated microbiome has only recently been examined [35,84]. Observational studies can identify associations between microbial communities and host condition but cannot readily determine whether microbial differences contribute to or result from host changes [85–87]. Experimental manipulation provides one approach for evaluating these relationships, and antibiotic-mediated depletion has been used across diverse host systems to examine host responses to altered microbial conditions [19–21,88,89]. Similar to the restructuring observed here, short-term exposure to filter-sterilized seawater in *Exaiptasia diaphana* reduced bacterial diversity and altered community composition without detectable effects on host health or asexual propagation [90].

Microbial treatment accounted for the largest differences in polyp microbiome composition, while temperature contributed additional structuring that was more apparent under MES conditions but modest relative to the separation between microbial treatments (Fig. 4; Fig. S6-7). The resulting bacterial communities differed most notably at the class level. ABS communities were dominated by Bacteroidia, largely represented (59-89%) by the genus *Muricauda*, whereas MES communities contained a more diverse Alphaproteobacterial assemblage distributed across eight orders, with *Ruegeria* representing the most abundant named genus (9-41%). *Ruegeria* has been detected in the surface microbiome of wild Florida Keys *C. xamachana* medusae and identified as a dominant core genus in laboratory polyps, particularly during the mother-polyp and early-strobila stages [83,84]. Members of *Ruegeria* have been associated with vitamin and cofactor provisioning, sulfur metabolism, antioxidant production, and nitrogen processing in marine holobionts [84,91]. In contrast, *Muricauda* has been associated with Symbiodiniaceae and proposed as a competitive or opportunistic colonizer under altered microbial conditions [90,92]. Coral-associated *Muricauda* isolates also possess genes associated with carotenoid antioxidant production and reactive nitrogen species processing [91]. These reported associations and functional capacities provide context for *Ruegeria* and *Muricauda* in the contrasting communities observed here, but genus-level identity and relative abundance cannot establish whether these functions were expressed or contributed to host phenotypes. Although our bacterial communities were shaped by the microbial reservoir of a closed laboratory system, their overlap with taxa reported in wild *C. xamachana* medusae, laboratory polyps, and cultured Symbiodiniaceae suggests that some community members occur across study systems.

Shifts in bacterial communities have also been linked to algal presence in the holobiont. In *Cassiopea*, bacterial community composition changes during establishment of algal symbiosis and following symbiont loss [35,93], and cultured Symbiodiniaceae experiments indicate that associated bacteria can alter algal physiology in temperature-dependent ways [94]. In our study, reduced bacterial representation in ABS samples was accompanied by a greater proportion of 16S reads classified as algal chloroplasts prior to their removal from the dataset (Fig. S5). Because these data are compositional, however, this could reflect reduced bacterial reads rather than increased algal abundance, which was not directly quantified. Considered alongside previous studies, this shift raises the possibility that bacterial and algal components contribute to holobiont stability, such that disruption of one may be accompanied by changes in the other. Representative images of polyps and ephyrae also showed conspicuous loss of pigmentation at 33°C under both microbial treatments (Fig. S8), consistent with disruption of the photosymbiotic state at the upper seasonal temperature regardless of bacterial community state. Together, these findings highlight the importance of considering the host, algal symbionts, and bacterial communities when evaluating holobiont responses to seasonal temperatures. Direct measurements of algal density and physiology alongside bacterial activity and function will be needed to resolve the mechanisms underlying these interactions.

### Conclusions

Together, this study demonstrates that temperature and microbial context jointly shape multiple fitness-related traits in *C. xamachana* polyps and ephyrae, but their roles are not equivalent. Temperature primarily determined developmental and reproductive trajectories, whereas microbial context modified the magnitude and outcome of these processes. Temperature effects differed among seasonal conditions, with 20°C supporting polyp growth and budding without strobilation, 28°C supporting sustained ephyra production, and 33°C accelerating initial ephyra release but ultimately truncating strobilation and yielding smaller ephyrae with poorer condition and survival. Although developmental and reproductive processes remained functional following substantial bacterial depletion and restructuring, divergence between microbial treatments at 33°C suggests that microbial context may become increasingly consequential near the upper seasonal extreme. By linking conditions experienced by benthic polyps to the production and condition of ephyrae, these findings suggest that climate-driven changes in the timing, duration, and frequency of seasonal temperatures could alter favorable reproductive windows and ultimately impact *Cassiopea* medusa population dynamics.

## Acknowledgements

We thank Dr. Amy Tan, Stephanie Hendricks, and Myles Wagner for their thoughtful comments and extensive feedback on the manuscript, and members of the Strader Lab for helpful discussions throughout the development of this work. We acknowledge Dr. Megan Maloney for providing the founding polyps and Ziyu Wang for his contributions to establishing the laboratory stock used for this study. Portions of this research were conducted with the advanced computing resources provided by Texas A&M High Performance Research Computing.

## Contributions

M.M.C. and M.E.S. conceived the study. M.M.C. performed the experiment, analyzed the data, and drafted the initial manuscript. M.E.S. provided funding, project supervision, and contributed critically to writing, editing, and approval of the final draft.

