## Supplementary figures for "Seasonal temperature extremes drive shifts in reproductive strategy in *Cassiopea xamachana*"

***Supplementary figure 1.*** Seasonal water temperature variation in the upper Florida Keys
***Supplementary figure 2.*** Additional alpha- and beta-diversity metrics for $T_{35}$ bacterial communities
***Supplementary figure 3.*** Changes in polyp abundance and abundance-normalized reproductive output
***Supplementary figure 4.*** Timing and magnitude of peak bud production
***Supplementary figure 5.*** Taxonomic classification of 16S sequencing reads prior to filtering
***Supplementary figure 6.*** Bacterial community composition across taxonomic ranks in $T_{35}$ polyp samples
***Supplementary figure 7.*** Genus-level composition of $T_{35}$ polyp bacterial communities
***Supplementary figure 8.*** Representative images of *Cassiopea xamachana* polyps and ephyrae

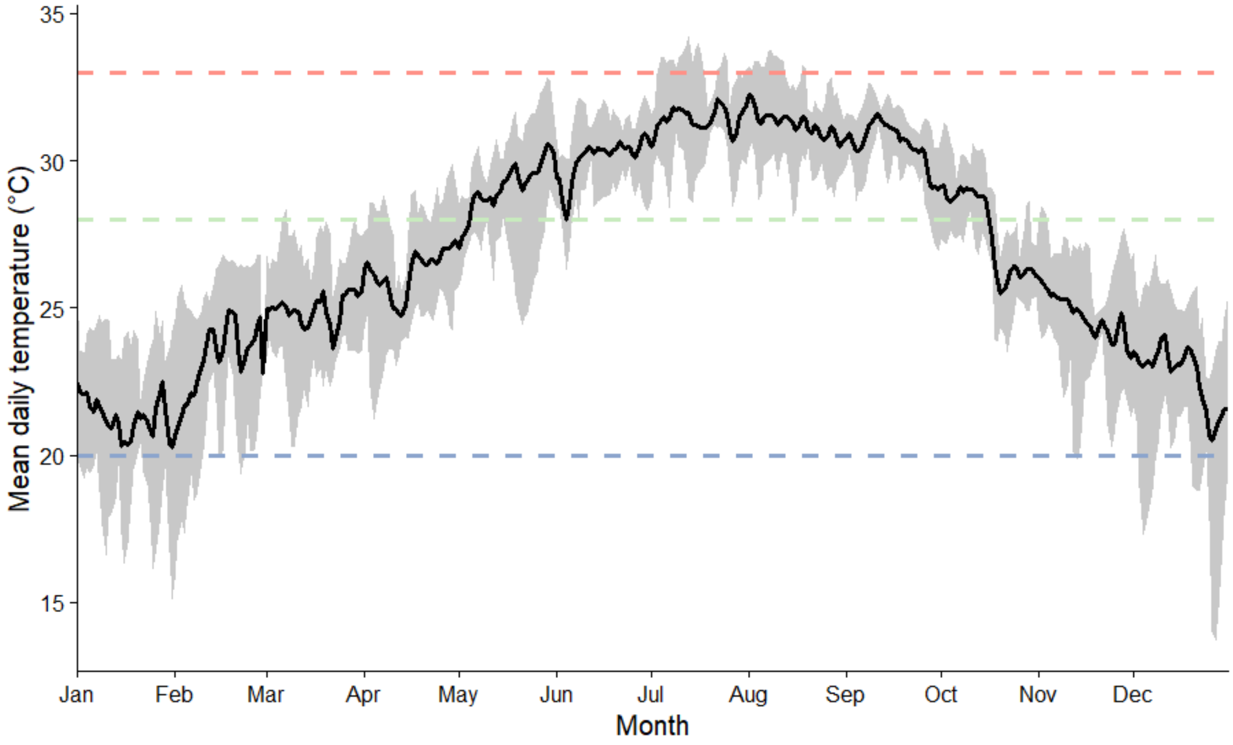


***Supplementary figure 1.*** Seasonal water temperature variation in the upper Florida Keys. Daily mean water temperatures from NOAA National Data Buoy Center (NDBC) station BNKF1 (25.087° N, 80.519° W) from 2021-2025. The black line represents mean daily temperature across the five-year period, and the grey shaded region represents the observed range of daily mean temperatures among years for each calendar day. Dashed horizontal lines indicate the experimental temperatures in this study: 20°C (winter seasonal extreme), 28°C (ambient temperature), and 33°C (summer seasonal extreme). Data source: NOAA National Data Buoy Center, Station BNKF1, Butternut Key, FL; accessed September 9, 2026.

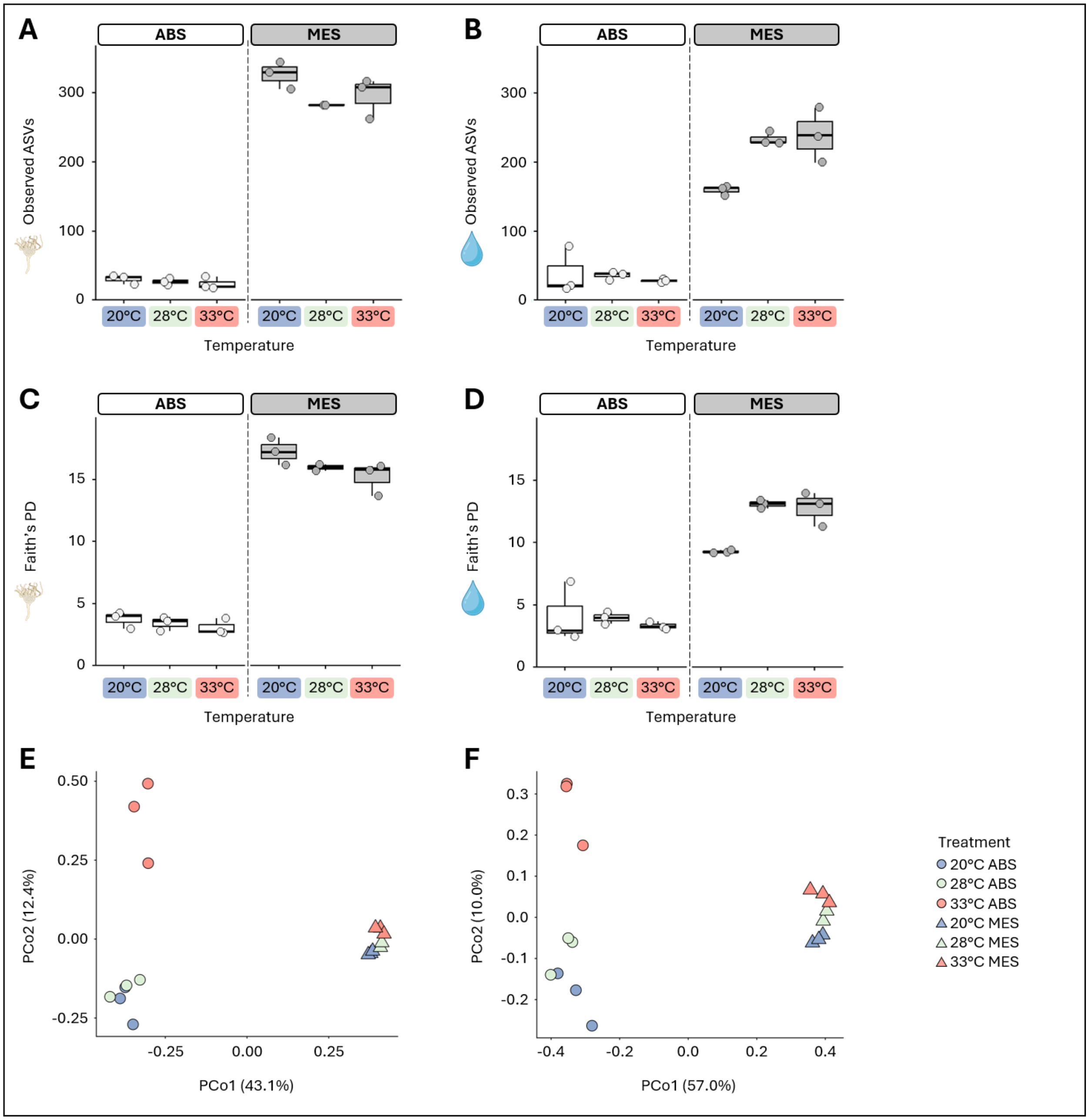


***Supplementary figure 2.*** Additional alpha- and beta-diversity metrics for $T_{35}$ bacterial communities. Boxplots of Observed ASV richness for polyp (A) and water (B) samples and of Faith's Phylogenetic Diversity (PD) for polyp (C) and water (D) samples. Boxes represent the median value and interquartile range (IQR), with whiskers extending to extremes within 1.5× the IQR. Points represent biological replicates. Principal Coordinates Analysis (PCoA) of $T_{35}$ polyp bacterial communities based on Jaccard dissimilarity (E) and Unweighted UniFrac (F) distance. Points are colored by temperature and shaped by microbial treatment.

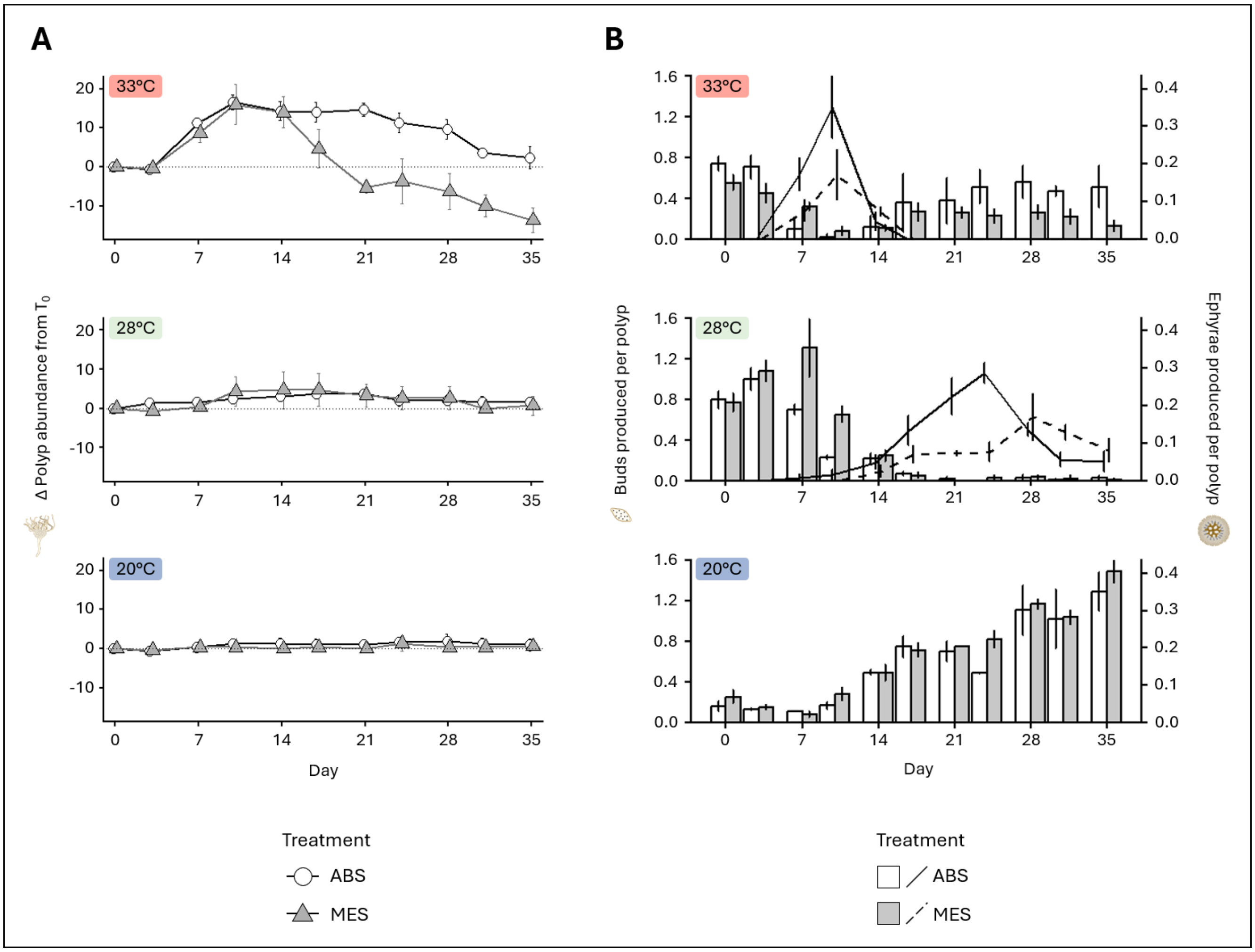


***Supplementary figure 3.*** Changes in polyp abundance and abundance-normalized reproductive output through time. Change in polyp abundance relative to $T_0$ across the 35-day experiment (A). Points represent mean change in polyp abundance ± standard error for ABS (circles) and MES (triangles), with the dotted horizontal line indicating no change from $T_0$. Mean bud (bars; left y-axis) and ephyra (lines; right y-axis) production per polyp at each census interval ± standard error are shown in (B). Offspring production was normalized by the number of polyps present at the beginning of each census interval. ABS is shown in white/solid lines and MES in gray/dashed lines.

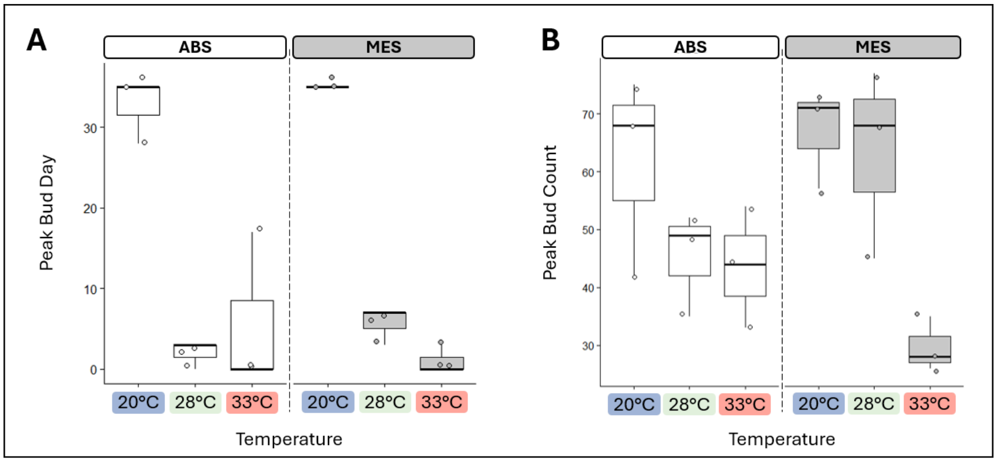


***Supplementary figure 4.*** Timing and magnitude of peak bud production. Experimental day on which peak bud production occurred (A) and maximum number of buds recorded at a single census (B) for each treatment. Boxes represent the median value and interquartile range (IQR), with whiskers extending to extremes within 1.5× the IQR. Points represent biological replicates.

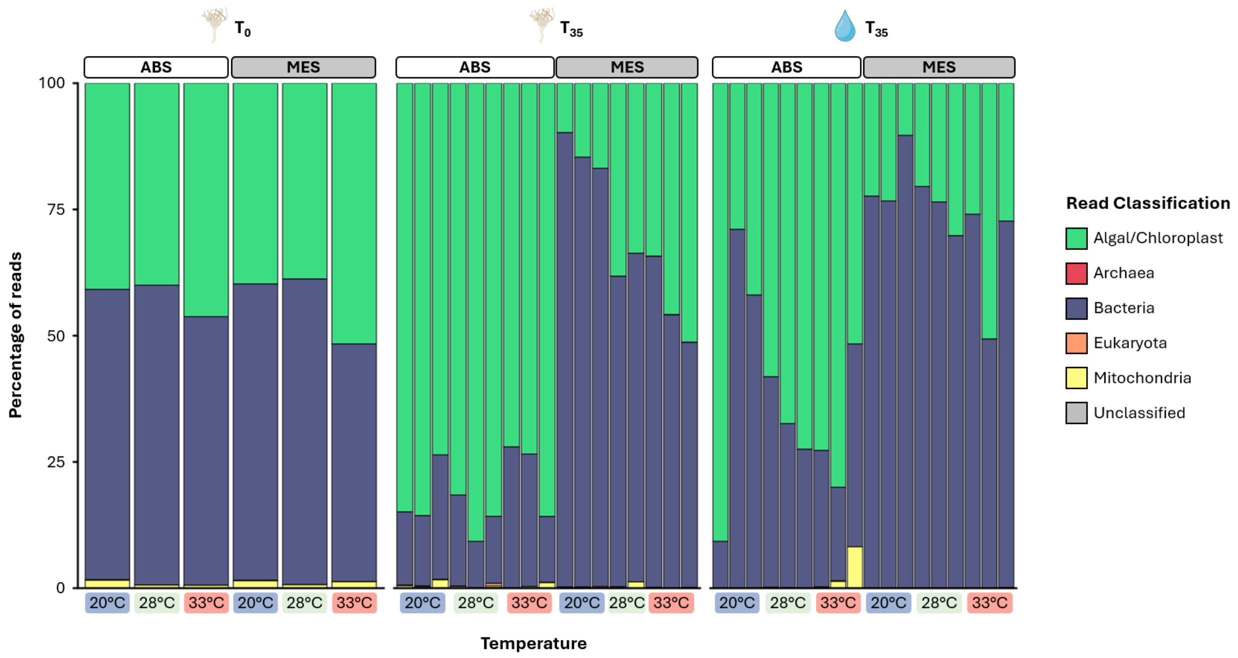


***Supplementary figure 5.*** Taxonomic classification of 16S sequencing reads prior to filtering. Percentage of reads assigned to each classification for $T_0$ polyp samples (left), $T_{35}$ polyp samples (middle), and $T_{35}$ water samples (right). Across samples, Algal/Chloroplast reads comprised 9.8-90.8%, Archaeal reads < 0.1%, Bacterial reads 9.2-90.0%, Eukaryotic reads < 0.5%, Mitochondrial reads < 8.2%, and Unclassified reads < 0.1%. Only bacterial reads were retained for downstream microbiome analyses.

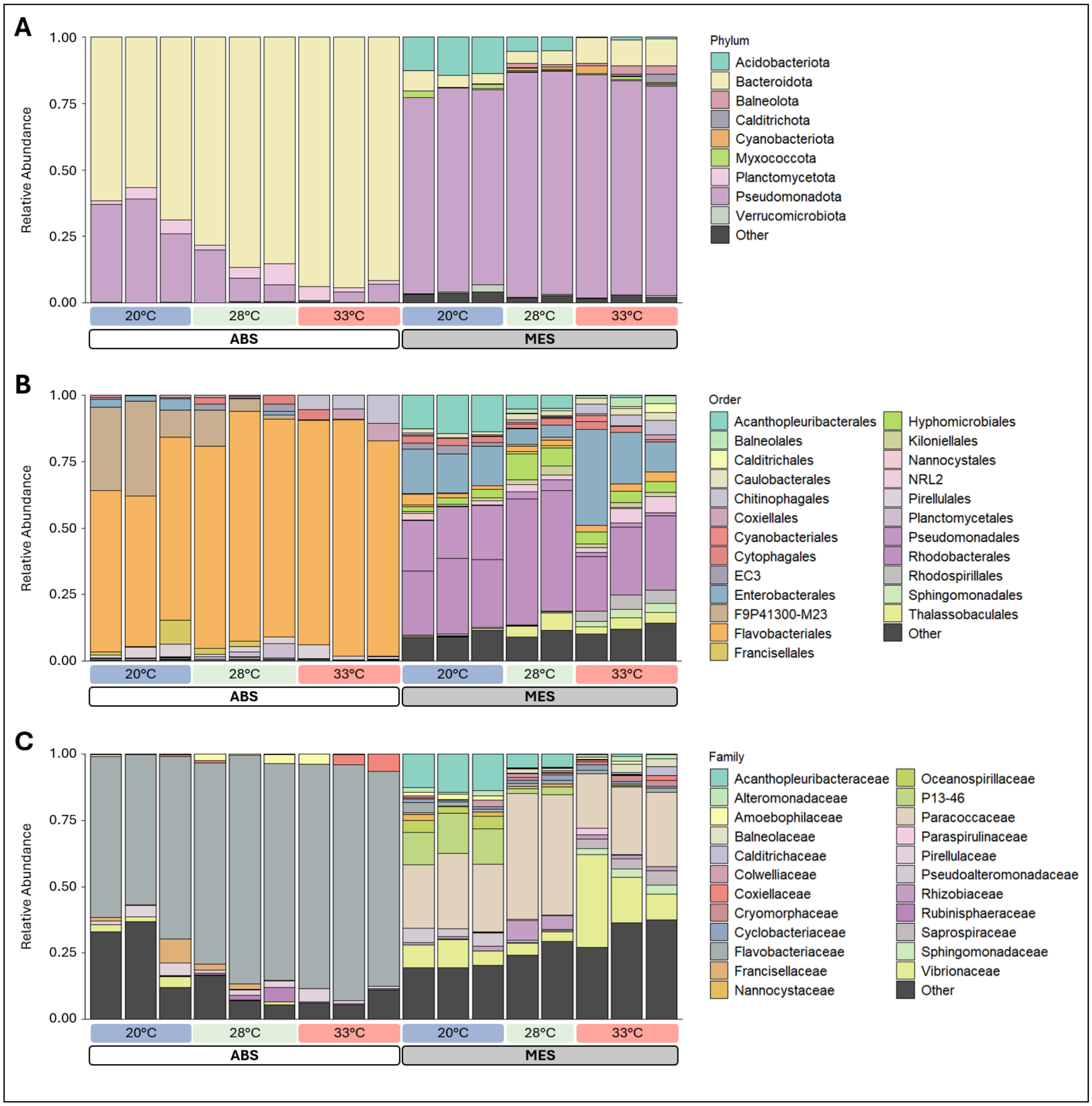


***Supplementary figure 6.*** Bacterial community composition across taxonomic ranks in $T_{35}$ polyp samples. Stacked barplots show the relative abundance of bacterial taxa at the phylum (A), order (B), and family (C) levels across temperature and microbial treatments. The "Other" category represents unclassified bacterial taxa and taxa below the 2% total abundance threshold.

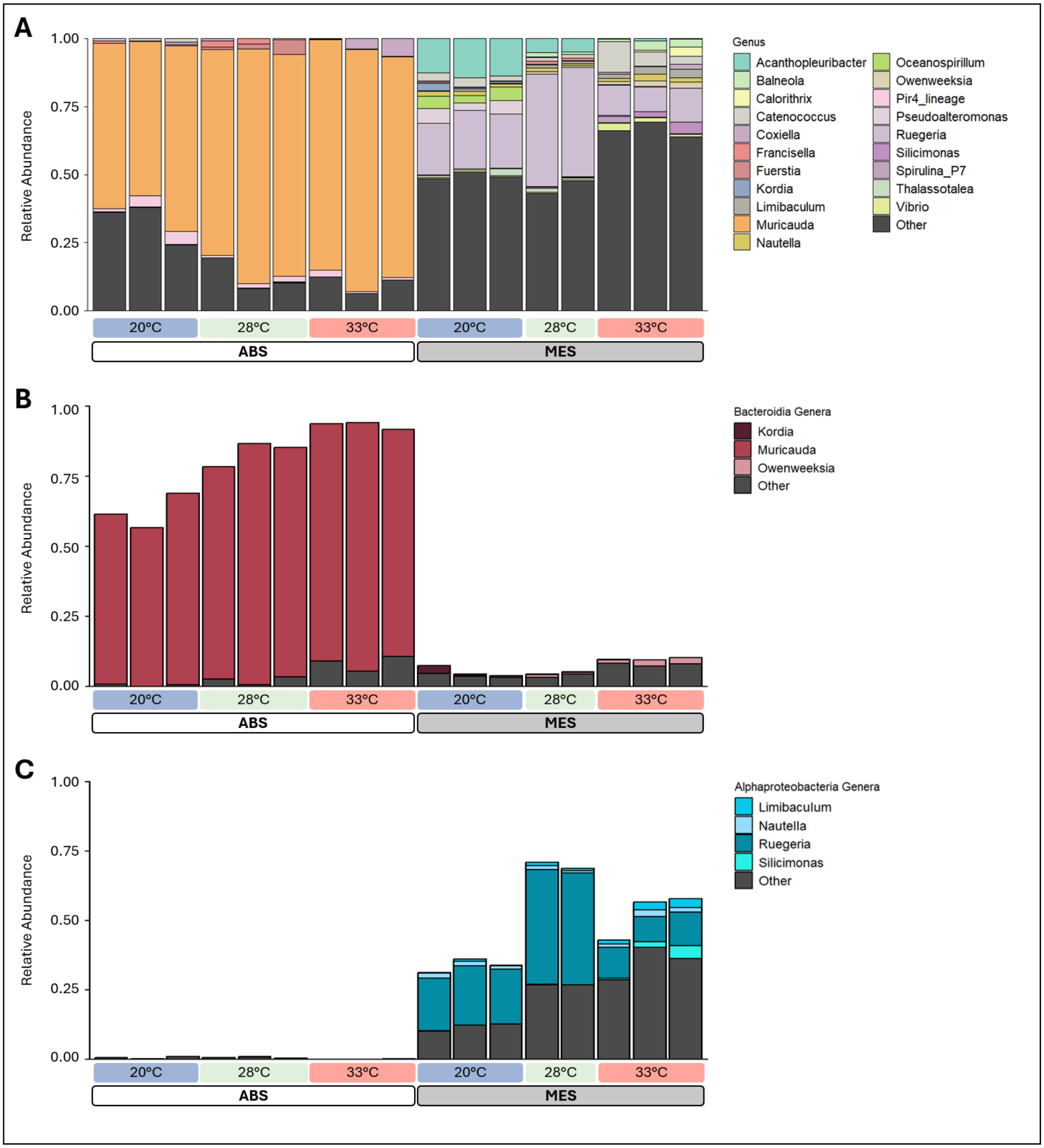


***Supplementary figure 7.*** Genus-level composition of $T_{35}$ polyp bacterial communities. Stacked barplots show genus-level relative abundance for the total bacterial community (A), for genera within Bacteroidia (B), and for genera within Alphaproteobacteria (C) across temperature and microbial treatments. Relative abundances in B and C were calculated from the total bacterial community and were not renormalized following class-level filtering. In A, the "Other" category represents unclassified bacterial taxa and genera below the 2% total abundance threshold. In B and C, "Other" represents genera unclassified within the respective class and named genera below the 2% total abundance threshold.

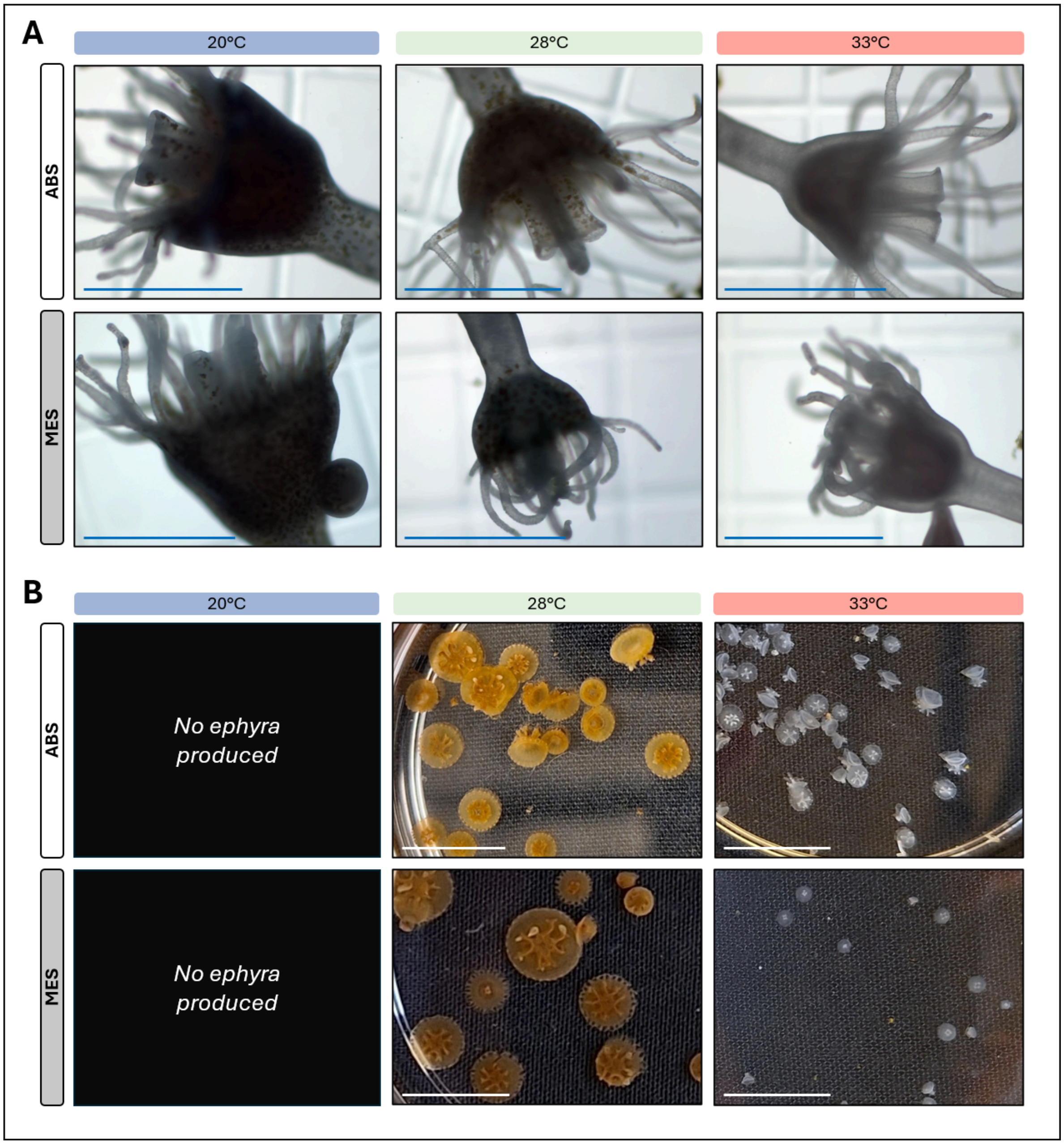


***Supplementary figure 8.*** Representative images of *Cassiopea xamachana* polyps and ephyrae across temperature and microbial treatments. Representative polyps (A) photographed on Day 34 of the 35-day experiment. Representative ephyrae (B) photographed at the final census (Day 35). No ephyrae were produced at 20°C. Scale bars = 1mm (A) and 1cm (B).
